# ACI-1 class A beta-lactamase is widespread across human gut microbiomes due to transposons harboured by tailed prophages

**DOI:** 10.1101/235788

**Authors:** Chris M Rands, Elizaveta V Starikova, Harald Brüssow, Evgenia V Kriventseva, Vadim M Govorun, Evgeny M Zdobnov

## Abstract

Antibiotic resistance is increasing among pathogens at unprecedented rates and the human body contains a large pool of antibiotic resistance genes that can be spread among bacteria by mobile genetic elements. *Acidaminococcus intestini*, a bacterium found in the human gut that belongs to the class of Negativicutes, is the first gram-negative coccus shown to be resistant to beta-lactam antibiotics. Resistance is conferred by *aci1*, a gene encoding the ACI-1 class A beta-lactamase, but the evolutionary history of *aci1* and its distribution across other Negativicutes and in the human gut microbiota remains obscure. We discovered that ACI-1 proteins are phylogenetically distinct from class A beta-lactamases of gram-positive Firmicutes and that the *aci1* gene occurs in bacteria scattered across the Negativicutes clade, suggesting possible mobilization. In the reference *A. intestini* RyC-MR95 strain, we found that *aci1* is surrounded by mobile DNA, transposon derived sequences directly flank *aci1* and are likely the vehicle for its mobility. These transposon sequences reside within a prophage context consisting of two likely degraded tailed prophages, the first prophages to be characterised in *A. intestini.* We found *aci1* in at least 56 (4.4%) out of 1,267 human gut metagenome samples, mostly hosted within *A. intestini*, and, where could be determined, mostly within a similar constellation of mobile elements to that found in the reference *A. intestini* genome. These human samples are from individuals in Europe, China and the USA, showing that *aci1* is widely distributed globally. Additionally, we examined the nine different Negativicute genome assemblies that contain *aci1*, and found that only two of these strains show a similar mobile element context around *aci1* to the reference *A. intestini* with transposons adjacent to a tailed prophage. However, in all nine cases *aci1* is flanked by transposon derived sequences, and these sequences are diverse, suggesting the activity and degradation of multiple transposons. Overall, we show that ACI-1 proteins form a distinct class A beta lactamase family, and that the *aci1* gene is present in human guts worldwide within Negativicute bacterial hosts, due to transposons, sometimes inserted into tailed prophages.

## Introduction

Antibiotic resistance is one of the most pressing global health issues as hospital and community acquired infections are acquiring resistance at high rates [1–3], while the dearth of novel antibiotic classes makes this particularly concerning [4]. Bacterial genes undergo horizontal gene transfer frequently [5] and antibiotic resistance gene (ARG) transfer is a major mechanism by which resistance spreads [6]. ARG surveys have historically focused on pathogens from classical gram-positive and gram-negative bacteria. Numerous different mobile elements types have been implicated in spreading ARGs within pathogens, most frequently plasmids and transposons [6,7]. Interestingly, prophages have less often been proposed as major mobilizers of ARGs in pathogens, with only a few examples [8,9] of ARGs encoded within sequenced genomes from isolated phages [10,11]. By contrast, phages have long been known to encode important bacterial toxins and virulence factors [12,13], such as the Shiga, Cholera, diphtheria, and botulinum toxins [14–17].

However, the major reservoir of ARGs may not be pathogens but instead commensal bacteria residing in the bodies of humans and other animals, which can donate their ARGs to opportunistic pathogens [18]. Shotgun metagenomic sequencing has permitted the identification of thousands of ARGs in human gut microbiomes [19,20], with many being associated with mobile genetic elements [21], including phages [22]. Furthermore, antibiotic treatment of mice led to an increase in phage-encoded ARGs in metagenomes [23]. Nonetheless, the importance of phages in spreading ARGs within human and mice microbiomes remains disputed [11], and is important to investigate further, particularly given the renewed interest in using phages to target bacterial infections via phage therapy [24].

Some oral, vaginal and particularly intestinal, bacteria belong to the Negativicute class, a recently classified peculiar side branch of Firmicutes [25,26]. Firmicutes are generally low GC content gram-positive monoderms (with a single membrane), but the Negativicutes are diderms since they possess an outer membrane with lipopolysaccharides like gram-negative Proteobacteria. Little is known about mobile DNA in Negativicutes, for example, phages are known to exist in *Veillonella* [27] and *Selenomonas* [28], but nothing is known for phages of *Acidoaminococcus, Megasphaera*, and *Dialister*, important gut and vaginal bacteria. The characteristics of their phages would be interesting phylogenetically since they should display a hybrid character of phages obliged to cross outer membranes like phages from Proteobacteria and to function in a Firmicute-like cytoplasm. Furthermore, since they comprise widespread human commensal bacteria, it is medically important to test whether Negativicutes could be a source of ARGs and whether, and what class of, their mobile elements could spread these genes, potentially to both gram-negative and gram-positive bacteria.

*Acidaminococcus intestini* i s a Negativicute that forms part of the normal human gut microbiota but has also been found in clinical samples [29] and has recently been associated with polymicrobial infections and complex diseases, such as rosacea, a chronic inflammatory dermatosis [30], and growth faltering [31]. Importantly, *A. intestini* is the first gram-negative coccus with demonstrated resistance to beta-lactam antibiotics. The reference *A. intestini* strain RyC-MR95 genome, which was sequenced from a perianal abscess sample isolated from a European male diabetic patient, contains the *aci1* gene, which encodes the ACI-1 class A beta-lactamase that confers resistance to penicillins and extended-spectrum cephalosporins [32]. It has been speculated that a transposon might be involved in the mobilization of *aci1* based on the GC content of the region in *A. intestini* [32], but no detailed analysis was conducted. More recently the whole *A. intestini* genome was sequenced [33] and that of another related strain [34], but the evolutionary history of *aci1* and its genomic context in *A. intestini* and other Negativicutes remains unclear. Furthermore, the frequency of *aci1* within the gut microbiome of the human population has not yet been quantified.

Here we investigated the phylogenetic position of the ACI-1 protein among other class A beta-lactamases, the mobile element context of the *aci1* gene in *A. intestini* and other Negativicutes, and its prevalence across human gut microbiomes. We found that the ACI-1 protein family is separate from other class A beta-lactamases of gram-positive Firmicutes, and that the *aci1* gene is surrounded by transposon derived sequences that likely mobilized the gene within Negativicutes. These transposons, are sometimes, such as in the case of *A. instestini*, inserted within tailed prophages. Furthermore, we found *aci1* in human gut samples, mostly within A. *instestini*, from across three continents, showing that the gene is widespread.

## Results

### ACI-1 proteins are distinct from other class A beta-lactamases and found in selected Negativicutes

*Acidaminococcus intestini* RyC-MR95 contains a functional copy of the *aci1* gene encoding the ACI-1 beta-lactamase class A resistance protein [32]. We found that the ACI-1 family of proteins is phylogenetically distinct from other class A beta-lactamases (**Figure 1**). A protein from *Mitsuokella jalaludinii*, a Negativicute, shares 57% amino acid identity to ACI-1 proteins and other homologous proteins, mostly from *Clostridium* and secondarily *Bacillus* (classical gram-positive Firmicutes), share at most 55% identity to ACI-1 and form clearly separate protein families.

**Figure 1:**
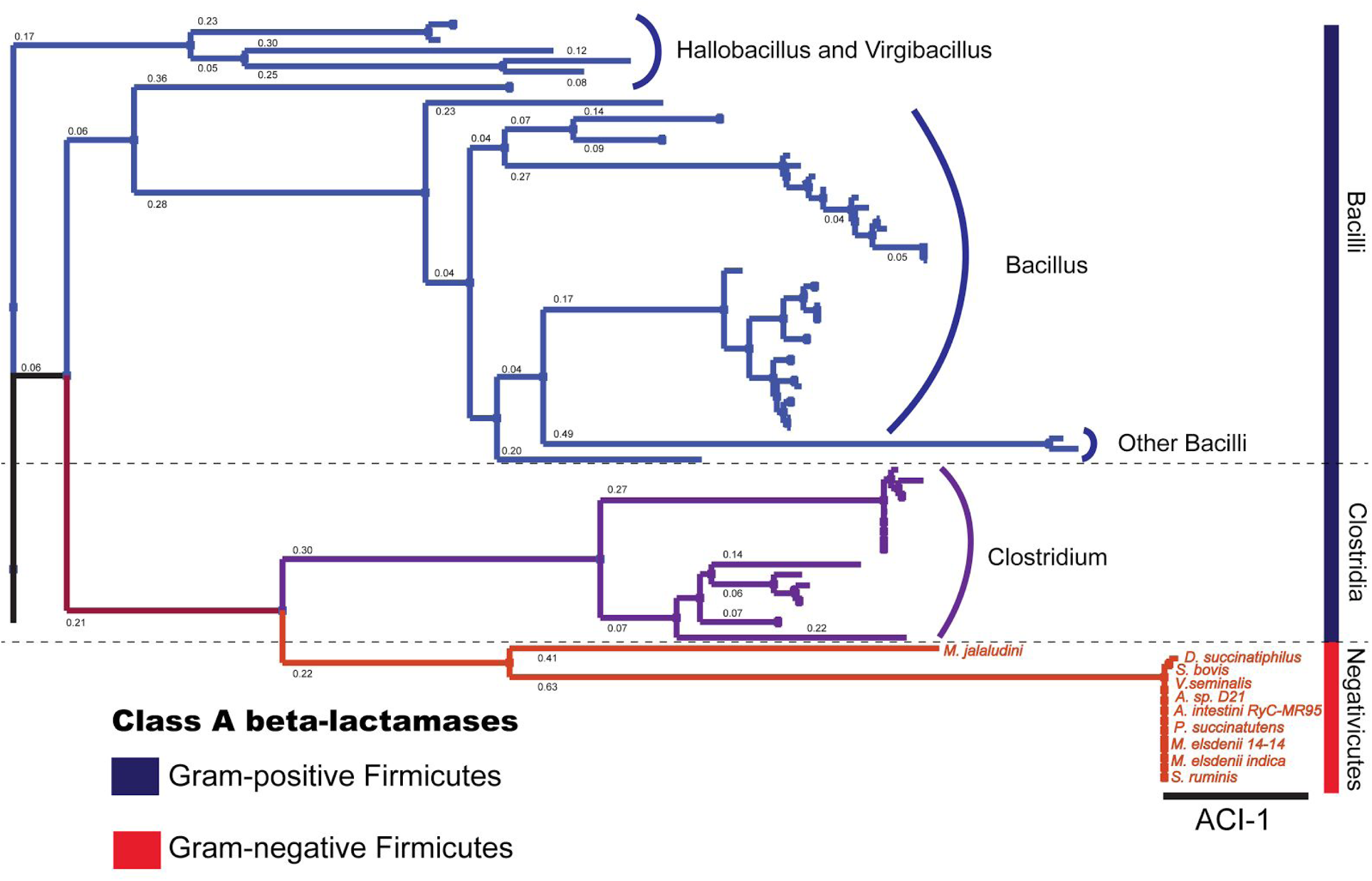
A phylogenetic tree of class A beta-lactamases. We found that ACI-1 proteins form a protein family separate from the other beta-lactamases that occur in gram-positive Firmicutes from Clostridia and Bacilli. The branch lengths with the amino acid substitution rates are shown; branch lengths of <0.04 are omitted to keep the numbers readable.

We found identical or near-identical copies of the *aci1* gene protein coding sequences in nine Negativicute genomes and notably its presence is phylogenetically patchy and not confined to a particular clade (**Supplementary Figure 1**). For example, we identified *aci1* in *A. intestini* RyC-MR95, *Selenomonas bovis*, and *Veillonella seminalis*, but not in complete genome assemblies from *Acidaminococcus fermetans* DSM, *Selenomonas sputigena* and *Veillonella parvula*.

### *aci1* is surrounded by transposons within a tailed prophage context in *A. intestini* RyC-MR95

Since the high sequence similarity of the identified ACI-1 proteins and their non-systematic presence in Negativicutes suggest acquisition by lateral and not vertical gene transfer, we looked for possible mobile vectors of the *aci1* gene in *A. instestini.* We found that *aci1* resides in a patchwork of different mobile elements, a set of transposon-derived sequences and two prophages (**Figure 2**). Distinct drops in the GC content between the mobile elements might be indicative of integration events, since the GC content is expected to be relatively homogenous across a prophage but can vary between a prophage, the lysogenic conversion module, and the bacterial host [35].

**Figure 2 :**
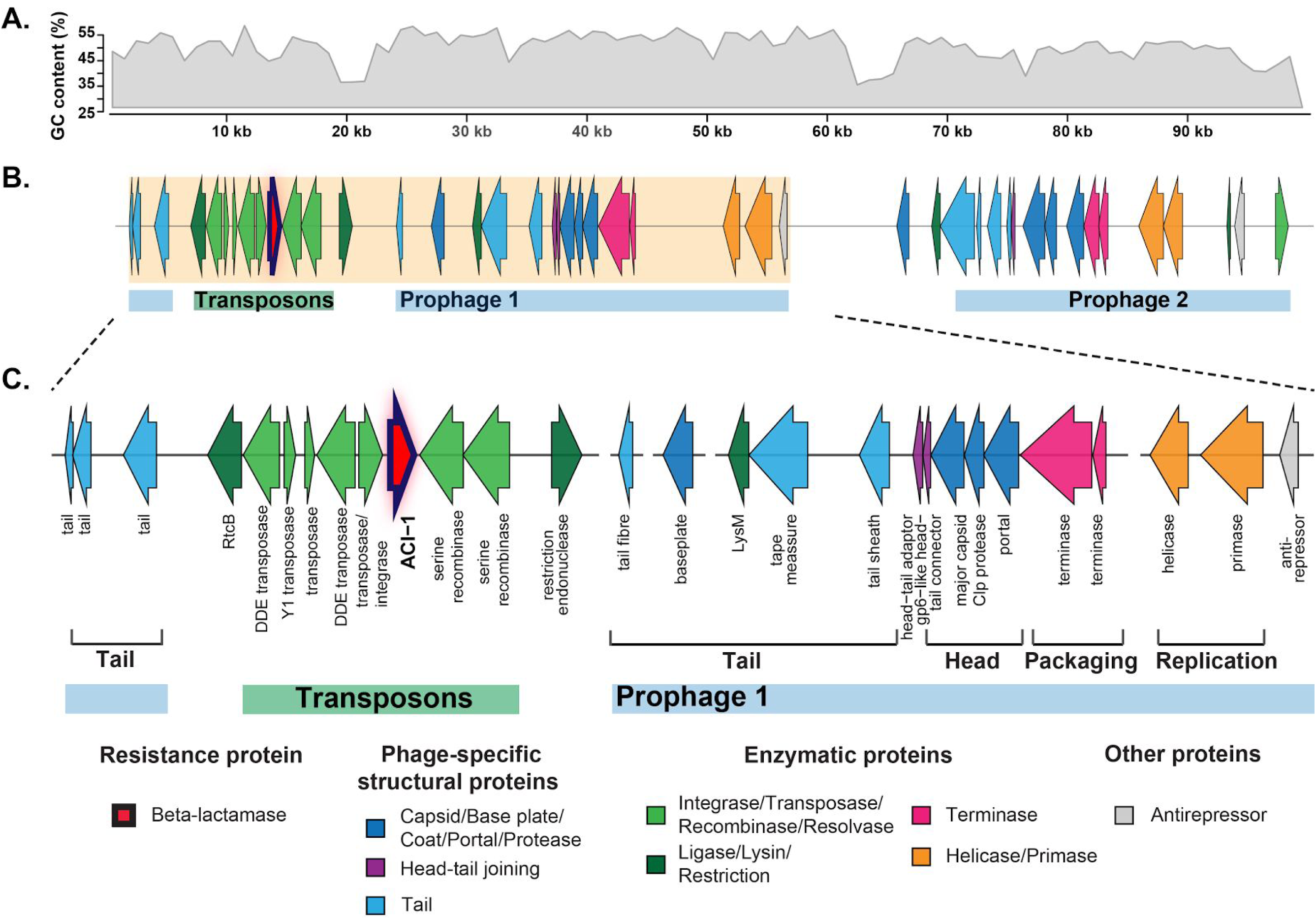
The genomic context of the *aci1* gene in the *Acidaminococcus intestini* RyC-MR95 reference situated next to a transposons and two prophages. We found that transposon derived sequences have likely carried *aci1* and integrated directly into a prophage. **A.** The GC content distribution for the region; the GC content dips around the boundaries of the mobile elements and might be indicative of integration events. **B.** The annotated proteins surrounding *aci1* with the transposon derived region and two prophages, and **C.** a more detailed view of the mobile elements nearest to *aci1* with annotated transposon related genes and Prophage 1, a novel tailed prophage that includes packing, head, head-tail joining, and tail units, like a classical Firmicute *Caudovirales*. The prophage is likely non-inducible as indicated by the fragmented tail structure and gap between the packaging and replication units. The 100kb region is shown to scale, each arrow is a protein with the arrowhead direction indicating the strand. Annotations for Prophage 2 are given in **Supplementary Figure 2**.

Since the transposon derived sequences directly flank *aci1*, they are most likely vehicle for its acquisition. These sequences consist of several different genes encoding enzymatic proteins adjacent to *aci1* that are of multiple origins: a DDE superfamily transposase, an IS200/IS605-specific Y1 type transposase, a transposase of unclear type, a DDE superfamily transposase possibly belonging to IS4 family, an integrase/transposase of unclear type, and two serine recombinases (**Figure 2**). We were not able to identify terminal repeats in this region, which are normally required for the active function of insertion sequences, but are not necessary for those of the IS200/IS605 family [36].

The transposons are located within a cryptic prophage setting (**Figure 2**). The context is prophage because the gene constellation from terminase to tail fibre resembles the classic *Caudovirales* (double-stranded DNA tailed prophages) structure previously observed in bacteria, including gram-positive Firmicutes [35,37–41]. Specifically, we annotated tail proteins, a head-tail joining region consisting of a head-tail adaptor followed by a gp-6 like head-tail connector protein. We found the head unit follows with a major capsid protein, Clp protease, and portal protein. Then the packaging unit comprises of large and small terminase subunits.

However, Prophage 1 does not display the standard genome organization for an inducible prophage. There is a gap between the packaging and replication unit. Additionally, the tape measure gene is too short for this class of Firmicute-like *Caudovirales,* the tail region is fragmented and disrupted by the transposons, and we found no clear lysin cassette (with a holin) and no repressors or antirepressors at the tail end that characterize a lysogeny module. Also the adjacent Prophage 2 represents only part of the standard genome for prophages from either Firmicutes or Proteobacteria. Its genome map includes identifiable tail, head-tail joining, head, packaging, replication, and lysogeny units (**Supplementary Figure 2**). However, Prophage 2 is too far from *aci1* to have been involved in its mobilization and also appears to be degraded, since it is missing several head or head-tail proteins and there is also a gap between the packaging and replication units.

These observations point towards two ancient prophages in the vicinity of *aci1* in *A. intestini* RyC-MR95, with Prophage 1 harbouring multiple more recent integration events from transposons, likely including an event that led to the acquisition of *aci1*.

### *aci1* is widespread across human gut microbiomes, mostly in a consistent constellation of mobile elements in *A. intestini*

Since *A. intestini* is a known human gut bacterium we investigated the prevalence of the *aci1* gene in human gut metagenome samples. Classifying reads to the bacterial class level using clade-specific taxonomic markers, we found at least one Negativicute strain in 1,229 (97.0%) of 1,267 gut samples, and, separately scanning the scaffolds, we found *aci1* in 56/1,267 (4.4%) of the samples (**Figure 3, Supplementary Table 1**). *aci1* was found in all examined human populations, from Europe, USA and China, and was not significantly more prevalent in any particularly population (**Supplementary Table 2**), showing that *aci1* is widespread but not ubiquitous across human guts.

**Figure 3 :**
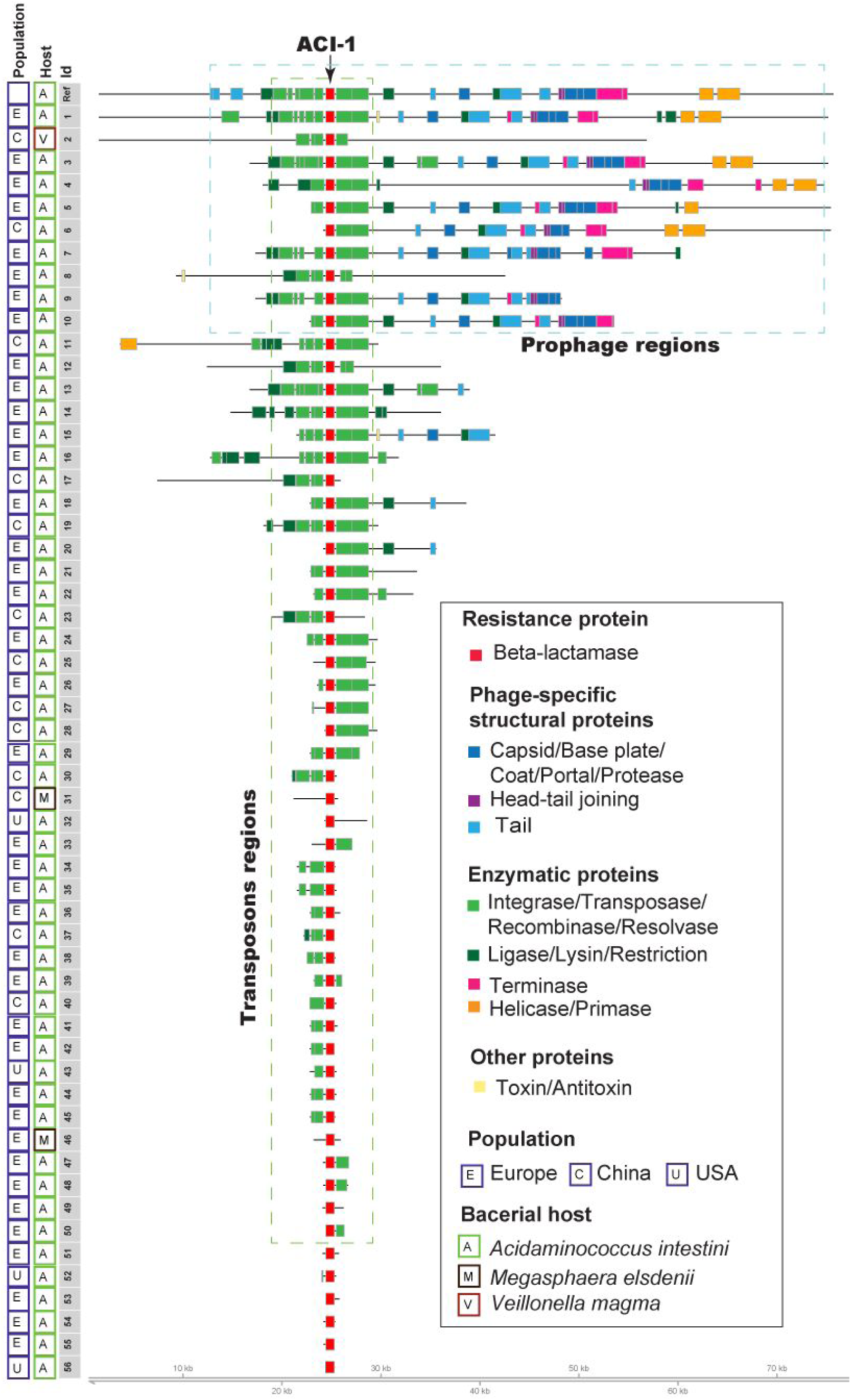
The prevalence and mobile element context of the *aci1* gene across *Acidaminococcus intestini, Megasphaera elsdenii*, and *Veillonella magna* from 56 human gut metagenomes. In all the populations from Europe, China and the USA, we identified copies of *aci1*. The majority of the scaffolds are from *A. intestini* and the larger scaffolds show a similar constellation of mobile elements around *aci1* that includes Prophage 1 and transposon derived sequences. The corresponding assembly and scaffold names are given in **Supplementary Table 1**. The protein level ‘alignments’ are anchored around *aci1* and black line behind the proteins shows the boundaries of the scaffolds.

We found the vast majority (53/56) cases with *aci1* within scaffolds that best aligned to *A. intestini* RyC-MR95. *A. intestini* scaffolds show a similar constellation of *aci1* surrounded by multiple transposon-associated recombination genes which, like a Russian doll, are sitting in a prophage as far as can be derived from the often short scaffolds. This suggests that where sufficient sequence is available, *aci1* is surrounded by transposons, and only secondarily by tailed prophage sequences.

Examining the remaining three cases of *aci1* found in the gut metagenomes that were not hosted within *A. intestini*, two were from short uninformative scaffolds from *Megasphaera elsdenii 14-14,* a Negavaticute mostly found in the rumen but also known to occur within the human gut [42,43]. The other case is from *Veillonella magna*, a Negativicute previously identified in swine intestines [44]. The metagenomic scaffold aligns almost perfectly to the reference *V. magna* scaffold with a single contiguous gap in the reference where *aci1* and surrounding transposon derived sequences reside (**Figure 4**), suggesting that this transposon unit alone, which is also present in the reference *A. intestini* strain, might be sufficient machinery to mobilize *aci1*.

**Figure 4:**
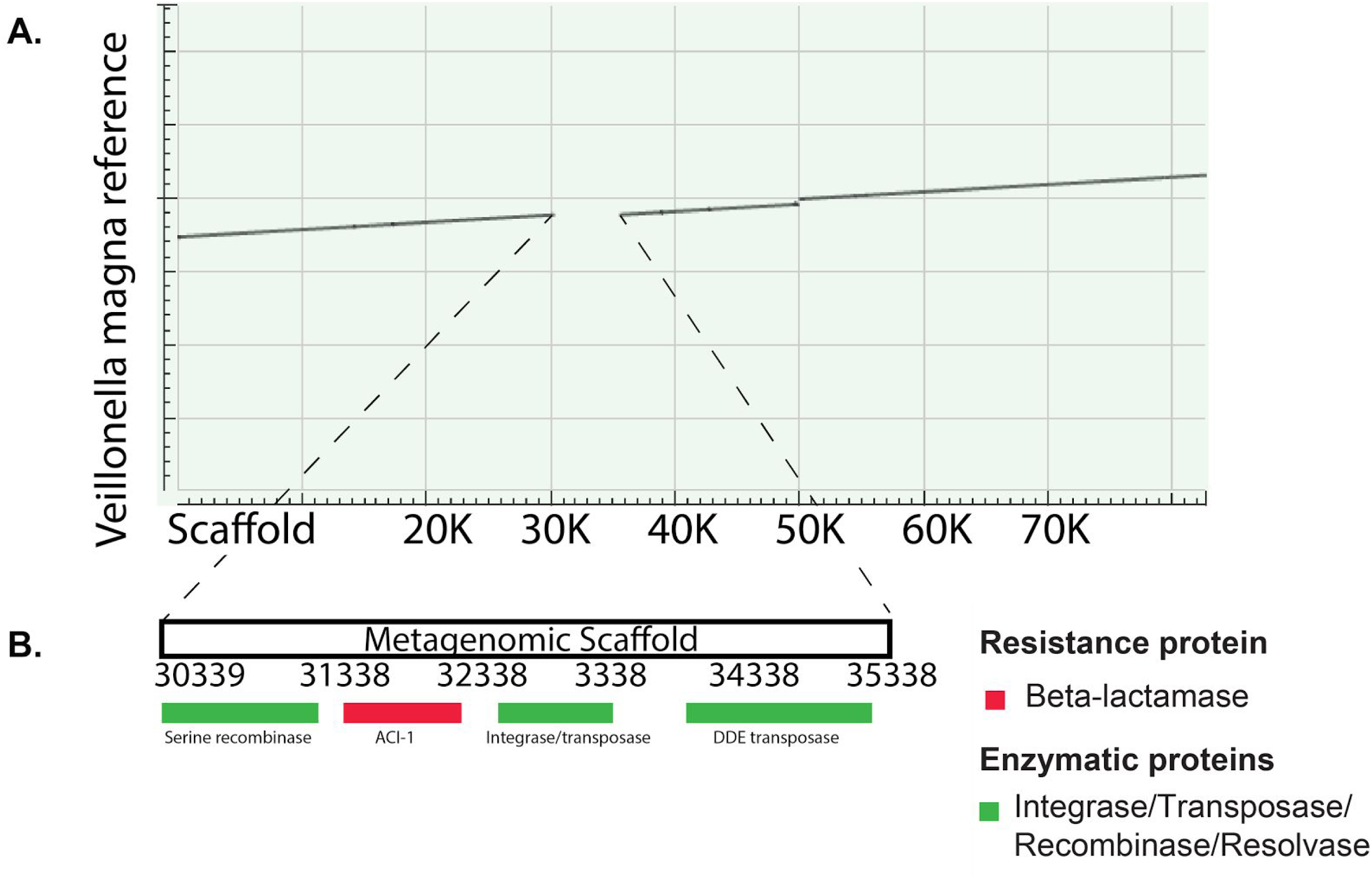
**A.** Dot plot alignment between scaffold32691_2 from the human gut metagenome assembly HT8A and the aligning *Veillonella magna* bacterial reference genome scaffold (Accession: AUAN01000002.1). The gap in the alignment is filled in **B.** where ACI-1 resides within several transposon annotated proteins. The absence of this transposon unit from the reference implies that it might be capable of autonomous mobilization.

### *aci1* is found in a diverse mobile element context across Negativicutes

We explored the mobile element context of *aci1* more widely across the Negativicutes. We built a phylogeny from the 16S ribosomal RNA sequences of the nine strains with *aci1* (**Figure 5A**), which shows as separate clades the Acidaminococcaceae, Veillonellaceae, and Selenomonadaceae families. Contrary to the near identical protein coding sequences of *aci1*, its surrounding mobile element context is diverse across the nine Negativicutes (**Figure 5B, Supplementary Table 3**). Only *A. sp. D21* and *M. elsdenii strain indica* show the same mobile element constellation as found in *A. instestini* RyC-MR95 with Prophage 1, while the *M. elsdenii 14-14* genome has a putative conjunctive element in the vicinity of *aci1*. A consistent trend across all strains, is that *aci1* is flanked by mobile element associated genes, such as transposases, recombinases, resolvases and integrases, but their number and orientation is variable. Further reinforcing this, when we performed a nucleotide alignment taking the genomic regions flanking *aci1*, we found that as the alignment extends from *aci1* in both directions, the regions do not align well across all nine genomes, indicating the activity and degradation of several different transposons (**Supplementary Figure 3**).

**Figure 5 :**
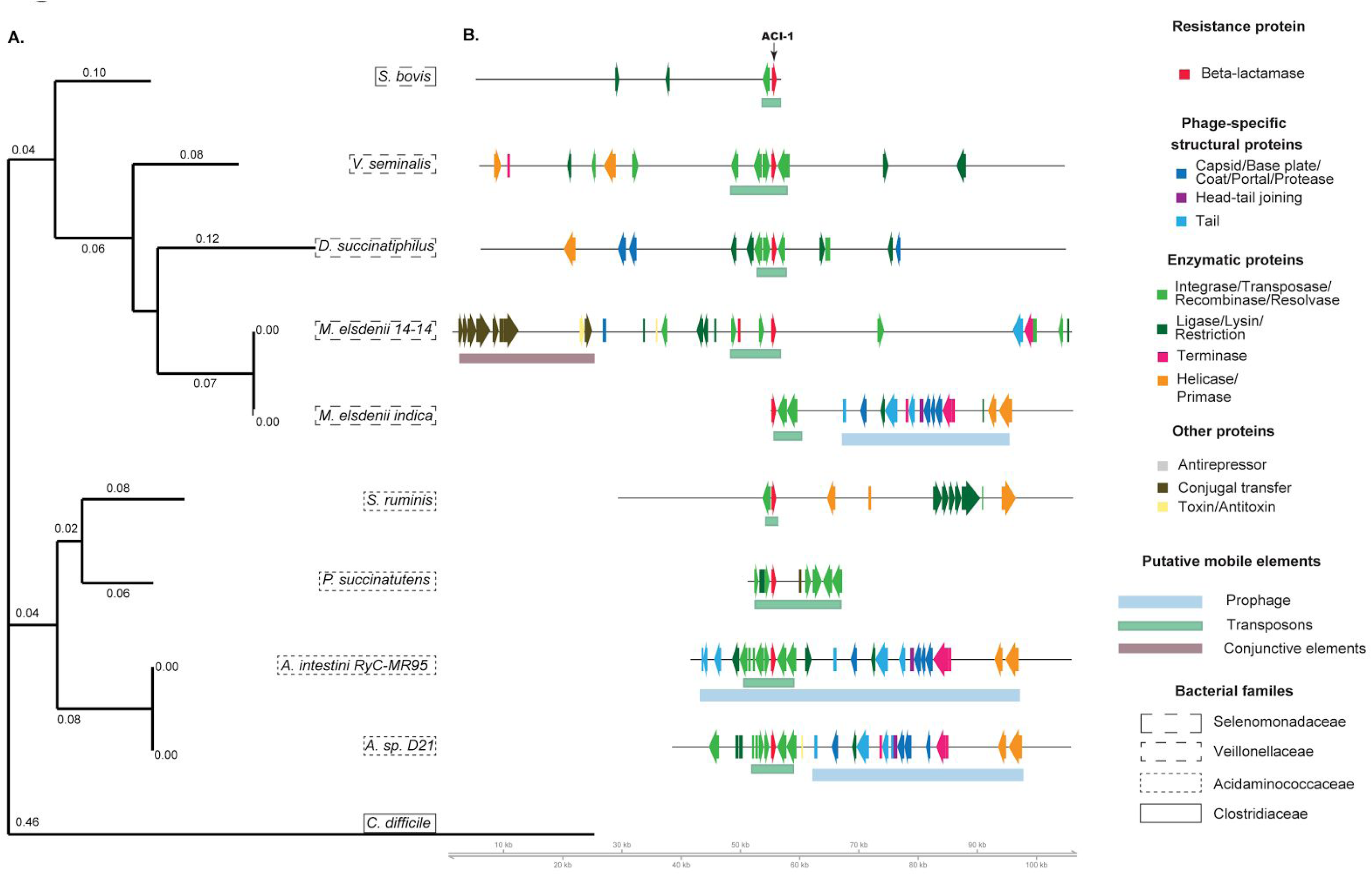
The species tree and mobile element context of *aci1* across nine Negativicutes. A. The species phylogeny is based on 16S ribosomal RNA sequences with the branch lengths to scale of the nucleotide substitution rate. *C. difficile* is the outgroup, a gram-positive Firmicute without *aci1*. **B.** the mobile element genes around *aci1* in each strain. We established that there is a diverse mobile element context surrounding *aci1* with two strains showing a similar pattern to that of the reference *Acidaminococcus intestini* strain, and remainder showing a different context without Prophage 1, but with diverse transposons. A nucleotide level alignment around *aci1* across the nine sequences is shown in **Supplementary Figure 3** and information on the assembly scaffolds is in **Supplementary Table 3**.

The diversity of mobile elements surrounding *aci1* in different Negativicute strains show that the evolutionary spread of *aci1* across Negativicutes is complex and likely involved multiple integration events from transposons, only occasionally within the prophage context we observed for *A. intestini*.

## Discussion

We found that *aci1*, a gene encoding the ACI-1 class A beta-lactamase that confers resistance to penicillins and expanded-spectrum cephalosporins [32], is surrounded by mobile DNA. Transposons, at least one of which likely carried *aci1*, have invaded a tailed prophage-rich region in *A. intestini* multiple times, suggesting that the region has characteristics attracting integration events. Proteobacteria T4-like phages are also targets of external mobile DNA [45], so phages from diverse bacterial hosts may facilitate the integration of other mobile element classes. When mobile elements co-locate like this, it will be non-trivial for systematic metagenome-wide studies to disentangle the roles of different mobile element types in transporting bacterial genes. Indeed, a less thorough investigation of this case might have concluded that prophages, rather than transposons, were the vehicle responsible for carrying *aci1*. This may partially explain the apparent paradox that isolated phage genomes rarely encode antibiotic resistance genes (ARGs), yet ARGs are frequently associated with phages in metagenomic and viromic studies [11].

Since *aci1* appears to reside in an integration hotspot in the genome, it is still at risk of further mobilization by transposons. This is of potential concern since transposons have been found in diverse bacteria [46] and might carry ARGs across wide phylogenetic distances [47], including to pathogens, because transposons often have recombinases that can operate in many different hosts without the need for host-specific factors. By contrast, the risk of future phage mobilization events seems low since prophages generally have a very narrow host range [48], and we found the prophage genome structure is not completely intact and therefore the prophages are likely not able to induce. These prophages are, to our knowledge, the first examples characterised in *Acidaminococcus* or *Megasphaera*. Strikingly, the prophages appears to follow the classical Firmicute *Caudovirales* genome structure, yet must have different machinery required to cross the additional outer membrane of these unusual gram-negative branch of Firmicutes. Although plasmids have been proposed as major mobilizers of ARGs, they do not appear to play a role in this case since we found no plasmid annotated proteins near *aci1*.

We found *aci1* is widespread across human gut microbiomes, being present in at least 4.4% of human gut microbiome samples we examined, including samples from Europe, USA and China. Previously, *aci1* has not been reported outside of Europe and the prevalence of it may be underestimated due to the limited and uneven coverage permitted by shotgun metagenomic sequencing. It has been proposed that ARGs in the human gut may pass between other animal guts [49], and since we found *aci1* in human gut bacteria normally found in animal bodies, *aci1* might be capable of undergoing cross-species transmission. The non-negligible frequency of *aci1* in the human gut microbiome and its presence across at least three continents is of medical relevance for at least two reasons, even when the gene is not directly transported into pathogens. First, *Acidaminococcus* and other Negativicutes have been found in clinical samples and may be involved in infections such as abscesses [29,50] or vaginosis [51] and have been associated with complex disorders [30,31]. Second, the secretion of beta-lactamases into the gut by commensal bacteria in response to beta-lactam antibiotics prescribed against bacterial pathogens could provide resistance for surrounding bacteria, including both pathogens and commensals, by collective resistance, sometimes termed indirect or passive resistance [52,53].

Our study shows that the *aci1* gene encoding a distinct family of class A beta-lactamase is present in the human gut microbiota across the globe, mostly within *A. intestini,* a common gut bacteria of unclear clinical significance. We find that *aci1* has likely been mobilized multiple times within the Negavaticitues due to transposons, sometimes harboured within tailed prophages. Investigation of other ARGs in this way would be valuable for understanding their prevalence and mechanisms of spread, particularly for genes from the understudied Negativicute clade, whose mobile genetic elements remain relatively uncharacterised despite the high prevalence of these bacteria in human bodies.

## Materials and Methods

### Genome and metagenome assemblies

We downloaded 160 Negativicute genome assemblies and accompanying protein annotations from NCBI Refseq [54], 18 of these assemblies are annotated as complete.

We obtained scafolds assembled from 1,267 human gut metagenome samples from an integrated catalog that includes data from the Human Microbiome Project, MetaHit and other projects [55].

### Identifying the *aci1* gene

We defined a copy of the ACI-1 protein as cases where the multispecies annotated ACI-1 protein (Accession: WP_006555379.1) had a BLAST hit with 99-100% amino acid identity and coverage and a copy of the *aci1* gene as cases where the *aci1* nucleotide sequence (Accession: AJ007350.1) had a 99-100% nucleotide identity and coverage hit. The ACI-1 protein covers 73% of the *aci1* gene nucleotide sequence, the exact internal translated sequence, but not the flanking regulatory sequences.

### Annotating sequences, genes, and mobile elements

To examine the mobile element context of a chromosome or scaffold with *aci1*, we predicted phage and phage-like regions using PHASTER [56]. Separately, we predicted proteins with MetaGeneMark version 3.38 [57] using default parameters for 50kb either side of *aci1*. Subsequently, we annotated proteins taking the top ranked (in terms of E-value) annotated BLASTp hit from NR and by scanning each protein with HMMSCAN [58] against a database of viral protein families represented as profile hidden markov models from the prokaryotic Viral Orthologous Groups [59]. We inspected all regions manually to resolve discrepant annotations and define the boundaries of the mobile elements. We defined the prophages using the phage-specific structural proteins from the last annotated tail gene to the last annotated replication unit gene. We defined the transposon derived sequences based on annotated transposases, recombinases, integrases and/or resolvases.

We carried out visualization of the gene models using the gviz R bioconductor package [60]. We did not display as arrows annotations for proteins that were not ARGs or virulence factors and were uninformative on the mobile element context, such as hypothetical proteins, but the regions are kept to scale.

We calculated GC content with Biopython [61] by dividing each sequence into 1kb windows and taking the mean GC content percentage for each window.

### Building alignments and phylogenies

For the ACI-1 protein phylogenies, we took the top (ranked by E-value) 100 hits from BLASTx of *aci1* gene nucleotide sequence against NCBI NR, excluding uncultured/unannotated sequences. After removing duplicates with identical fasta header descriptions, while ensuring all ACI-1 proteins were included, this left 72 beta-lactamase sequences. Note this excludes the beta-lactamase with accession AEQ22534.1 (which has 95% query coverage and 100% identity to the *aci1* nucleotide sequence) since the coordinates show it is actually a subset of WP_006555379.1 and not a separate protein, and protein with CCX56994.1, since we were unable to find an intact copy in the reference. Additionally we added a protein for *Megasphaera elsdenii* 14-14 that had a 99-100% nucleotide identity hit but not a protein hit, due to a frameshift or possible sequencing error, the frameshift was removed when calculating the ACI-1 protein phylogeny.

For the species phylogenies we took 16S ribosomal RNA (16S rRNA) sequences from the Ribosomal Database Project (RDP) [62] for each strain. For the phylogeny including taxa without *aci1*, we selected a single representative 16S rRNA for *Selenomonas sp. oral taxon* and for *Veillonella parvula* (for which multiple complete genome assemblies are available), and we excluded Negativicoccus massiliensis, since we were unable to find 16S rRNA sequences for this strain. For both species phylogenies we added in a *Clostridium difficile* sequence as an outgroup.

For both the ACI-1 and 16S rRNA sequences, we constructed multiple sequence alignments with MAFFT v7.309 [63] with a maximum of 1000 iterations. We then built Maximum Likelihood phylogenetic trees with iqtree v1.1.5 [64] using 1000 bootstraps and the TIM2 DNA model for the 16S rRNA sequences and the BLOSUM62 model for the ACI-1 proteins. We visualised the alignments and trees with Jalview [65].

For the nucleotide alignment around *aci1* across Negativicutes, we used MAFFT with a maximum of 1000 iterations and the --adjustdirectionaccurately flag.

### Negativicute abundances in the gut samples

We estimated the abundances of Negativicutes in each individual sample from the 1,267 human gut metagenomes using MetaPhlAn2 [66] applied to the raw reads from each sample.

### Enrichment test

To test for human populations with a significant enrichment for the *aci1* gene, we applied a hypergeometric test in R with a Bonferroni testing correction to account for the multiple populations tested. Specifically, for each population, we calculated a p-value that represents the probability that this number of *aci1* genes or more would occur in the population if *aci1* was distributed randomly across populations. Separately the fold change was calculated as (p_1_-p_2_)/p_2_; where p_1_ is the the proportion of samples from the population with *aci1* and p_2_ is the proportion of all samples with *aci1*.

## Acknowledgements

We thank Francisco Brito, Silas Kieser, Etienne Ruppé, Alexis Loetscher, and Andrey Letarov for useful discussions. We are grateful to Jennifer Tan for advice and help with making figures. We thank Nikita Prianichnikov for assistance with annotating phages.

